# A possible physical mechanism of the torque generation of the bacterial flagellar motor

**DOI:** 10.1101/2020.01.09.901124

**Authors:** Y. C. Chou

## Abstract

The torque required for the rotation of the rotor of a bacterial flagellar motor (BFM) can be generated from an impulsive force resulting from the collision between the stator and the rotor. The asymmetry in the fluctuations of the tilting angle of the rotor determines the direction of rotation. The expressions of the torque and the step size can be derived from a Langevin equation of motion. The drag coefficient of BFM derived from the Langevin equation and the measured torque–speed (τ-ω) relation is notably high; the viscous force from the environment cannot account for it. The drag force may be caused by the frictional interaction between the bearing-like L- and P-rings of BFM and the cell membrane. Order-of-magnitude estimations of the torque and the step size are consistent with previous experimental observations. The slope of the linear dependence of the rotational frequency on the temperature was estimated and was consistent with the observed value. A simulation device having the structural characteristics of BFM was designed to demonstrate the applicability of the proposed mechanism. Many observations for the actual BFM, such as the bidirectional rotation and the τ-ω relations of the clockwise and counterclockwise rotations, were reproduced in the simulation experiments.

**Importance:** The concept that the torque required for the rotation of the rotor of a bacterial flagellar motor (BFM) can be generated from an impulsive force resulting from the collision between the stator and the rotor is new and effective. The magnitude of the torque and the size of the step derived from the proposed mechanism are consistent with the observed values. The torque-speed (τ-ω) relation might be explained by the frequency-dependent drag force caused by the frictional interaction between the bearing-like L- and P-rings of BFM and the cell membrane. The slope of the linear dependence of the rotational frequency on the temperature is consistent with the observed value, which has not been achieved previously.

## Introduction

At any given moment, countless bacterial flagellar motors (BFMs) operate on earth. The basic mechanism that generates the torque required for the rotation of these sturdy nanometer-scale motors has remained unclear after five decades of intensive studies [1, 2]. The chemomechanical mechanism is the most popular theory [1, 3–6]. The general concepts of this mechanism are as follows: (a) the source of energy of the rotational motion is the electrostatic energy released by the ions (H^+^ or Na^+^) traveling through the electric potential difference across the cell membrane, (b) the stators gain the energy and perform a series of conformational changes (or power strokes), (c) the conformational changes may generate the torque required for rotor rotation, and (d) part of the rotational energy is released through filament motion in the viscous environment. Concepts (b) and (c) are the central subjects of studies attempting to understand the mechanisms of BFM. Many researchers have concentrated on solving the detailed structures of the stators and the rotor to understand the conversion from of electric energy to the mechanical energy of rotation [7–11]. The chemomechanical approaches had limited success in providing details of the torque-generating processes. When the ions travel along the designated path from the cell membrane to the stator, the electric potential energies of ions decrease and the kinetic energies increase. However, the excess kinetic energies of ions are released into the environment through collisions with the molecules in the path. Physically, the net effect of the ions traveling through the potential difference across the membrane is the increase in the kinetic energy of the random motion of the atoms and molecules along the traveled path. It is impossible to pass the entire gain in the kinetic energy of the ions to the stator in the form of chemical energy. Researchers have disagreed regarding the torque-generating processes; one consensus reached in the literature was that the torque was generated at the interface between the stator and the rotor [4–6, 9–12].

The rotation-related observations of BFM are summarized as follows. (a) The rotation is bidirectional and the direction of rotation can be switched instantaneously [2, 9, 13, 14]. (b) The torque-speed (τ-ω) relation indicates that the counterclockwise (CCW) and clockwise (CW) (viewed from the filament side) rotations are asymmetric [4, 5, 13, 15, 16]. (c) Steps in rotation are observed when the rotation is slow [17, 18]. (d) The number of active stators is not fixed [19–21]. The published literature contains controversies about whether the number of stators affect the rotational speed [6, 22]. A successful mechanism must explain or reproduce the aforementioned observations.

Mora et al. [5] and Mandadapu et al. [6] proposed a chemomechanical model of rotation by solving a set of Langevin equations for the stators and the rotor by including the conformation changes of the stators and the steric interaction potential between the stators and the rotor. I argue that the steric interaction potential that the authors used may not be the dominant interaction between the stators and the rotor. Although thermal fluctuations were included in the Langevin equations in the proposed chemomechanical model [5, 6], the effects of physical collisions between the stators and the rotor have been overlooked. The stators are in close contact with the outer lobe of the C-ring [6, 9, 23] of the rotor; however, the two bodies are not bound together and can move randomly and independently at room temperature. The amplitude of the thermal motion of the stator can be estimated with Δx ≈ (3k_B_T/k)^1/2^, where k_B_ is the Boltzmann constant, T is the environmental temperature, and k is the spring constant of the conformational change of the stator. Taking k in the order of 250 pN/nm and k_B_T = 4.1 pN.nm, Δx is of the order of 0.2 nm. If △x is larger than the gap between the stator and the rotor, frequent stochastic collisions between the two bodies are inevitable. The collisions may result in discontinuous changes in the trajectories of the two bodies involved. The impulsive force on the two colliding bodies is much larger than the force derived from a continuous interacting potential [5, 6]. Therefore, the impulsive force of the collision between the stator and the C-ring may be the dominant source for torque generation for the rotation of the rotor. The stators can be viewed as small studs that randomly collide with the C-ring.

Here, I propose a new physical mechanism that the impulsive force produced at the stator-rotor interface due to the random collisions between the two might be the dominant torque-generating force for the rotation of the rotor. The asymmetry in the direction of the tilt of the axis of rotation is vital for the rotation of BFM and may determine the direction of rotation. The rotation is bidirectional. The τ-ω relation depends on the direction of tilt and is found to be asymmetric for CCW and CW rotations. In the actual BFM experiments, the torque applied to BFM was deduced from the viscous force on an attached small bead. I would use the symbol τ_ext_ for the torque from the viscous force, because the viscous force is from the environment external to BFM [4, 15, 16], to distinguish τ_ext_ from the torque (τ_gen_) generated within BFM, which exists throughout the experiments for measuring the τ-ω relation. In the simulation experiments described later, τ_ext_ is applied by the gravitational forces of three hanging weights.

## Generation of impulsive torque

For the actual BFM, the asymmetry in the tilt of the rotor axis may result from the fact that the small bearings (L- and P-rings in actual BFM) may not be completely perpendicular to the outer membrane [24, 25]. The impulsive force, which is parallel to the stator axis, may not align with the tilted rotor axis. Therefore, a torque parallel to the rod (the rotor axis) may result. Asymmetry of a fraction of one degree in the tilting angle will be enough to generate processive rotations.

Fig. 1 shows the proposed mechanism for the generation of torque required for the directional rotation. The vertical axis is the apparent axis of rotation of the rotor. The rod corresponds to the rod of BFM and is the instantaneous rotation axis; the direction of the rod may fluctuate. The off-centered point P is the position where the stator collides with the rotor from above; therefore, the impulsive force 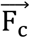 always points downward. The tilt of the rod is about the rotor axis with an angle α. In the figure, α is exaggerated. In the simulation experiments, α is within ±2°. Let ℓ be the projection of the vector 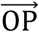 to the direction perpendicular to the direction of 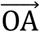 that asymmetry occurs, and is the lever arm of F_c_, where O is the center of the rotor (C-ring), and ϕ be the angle between the vector 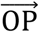 and the direction of 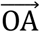. Then I = R sinϕ, where R is the distance between O and the stator. The impulsive force will exert a torque 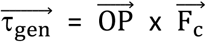 with respect to the rod. The magnitude τ_gen_ of the component of 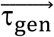 along the rod is

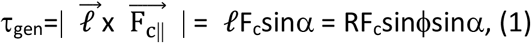

where 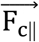 is the component of 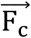 parallel to the plane of the rotor, and α is the azimuthal angle with respect to the motor (vertical) axis. The sign of τgen is positive when the rotation of the rotor is CCW. The magnitude of F_c_ can be estimated from a kinetic argument. The stator and the rotor are randomly moving and colliding with each other all the time. Assuming the relative speed of the rotor to the stator is v, then F_c_ = 2mv/△t, where m is the mass of the rotor, △t is the average time interval between two consecutive collisions, and 2mv is the change of momentum of the rotor in a collision with the stator. △t = 2d/v, where d is the spacing between the stator and the rotor. Therefore, the magnitude of 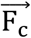 is F_c_ = mv^2^/d. In the one-dimensional motion, the average kinetic energy of the thermal motion of the rotor mv^2^/2 = k_B_T/2, where T is the temperature of the environment near the rotor. Thus, from Eq. (1)

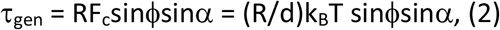

**Fig. 1.**
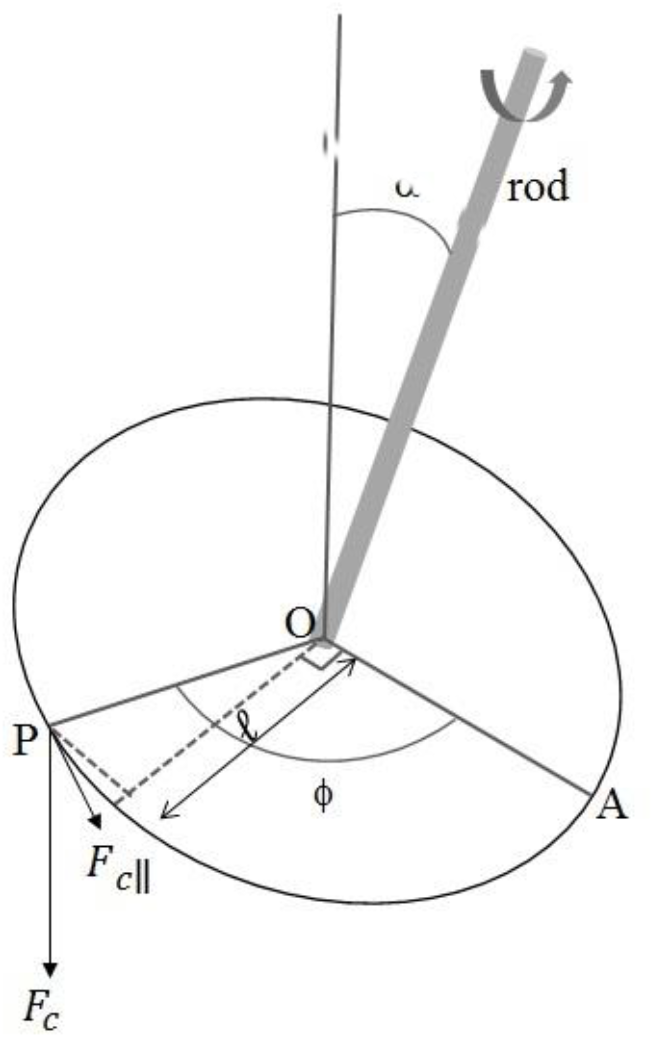
Schematic drawing of the mechanism for the generation of torque. The vertical axis is the axis of rotation. The tilted thick line represents the rod of BFM and is the instantaneous axis of rotation. The circle is the surface of the rotor, and point P is where the rotor and stator collide. The line OA is the direction of asymmetry in the fluctuations of the tilt angle α, and is the reference direction of the angle ϕ. The angle α can be <2°; in the figure, α is exaggerated. 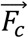 is the impulsive force on the rotor colliding with the stator. *F*_c||_ is the component of 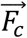 in the plane of the rotor. ℓ is the lever arm of *F*_*c*||_ about the rod.

Taking R = 20 nm, d = 0.2 nm, the thermal energy k_B_T = 4.1 pN.nm at room temperature, and sinϕsinα ≈ 0.1, τ_gen_ ≈ 41 pN.nm per collision.

The response of the rotor to the discrete impulsive torque can be formulated as a Langevin equation of motion. The equation of rotation can be written as

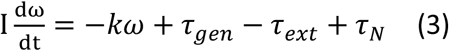

where I is the moment of inertia of the rotor, ω is the angular speed of the rotor, k is the drag coefficient of the rotor for the actual BFM or in the simulation experiments, τ_gen_= Σ_i_ τ_i_ *δ*(*t — t_i_*) is the torque on the rotor by the impulsive force, *τ_i_* is the ith impulse of torque occurs at *t_i_, δ*(*t - t_i_*) is the delta function which has non-zero value only within a time interval between *t_i_* and *t_i_* + Δ*t*, τ_ext_ is the constant externally applied torque (in the actual experiments [15, 16], τ_ext_ was from the viscous force on an attached bead), and τ_N_ is the ineffective contribution of torque from the thermal fluctuations of the rotor. We are interested in the long-time behavior of BFM (a time longer than the fastest time for one frame of image to be taken in the actual experiments ~ 0.1 ms [17, 18, 21]), during the time interval the average of *τ_N_* is zero.

After a change of variable, one solves Eq. (2) for the angular speed *ω_i_* for the ith impulse, and gets

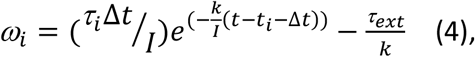

where τ_i_ and *τ*_i_Δ*t* are the torque generated and the impulsive angular momentum transferred to the rotor by the ith impulse, respectively, and Δ*t* is the time interval that the stator spends on one FliG in the C-ring. Supposing that BFM rotates at 200 Hz and 26 FliGs exist in the C-ring, one gets Δ*t* ≈ 2 × 10^-4^ s, which is notably close to the fastest time for the actual measurements up to the present. Within the time interval Δ*t*, the torques generated by different collision events with the same FliG might add up. Beyond Δ*t*, the collision events are not correlated, although the effects on the rotation of BFM might still add up. For simplicity, in this report I let Δ*t* = 1 × 10^-4^ s. The moment of inertial I of the rotor can be estimated from its mass m = 2.5 × 10^-20^ kg [1], by assuming the rotor being a cylinder with a radius of 20 nm, I ~ 5 × 10^-28^ kg.m. The value of the drag coefficient k can be obtained from the measured torque-speed (τ_ext_-ω) relation of the actual BFM; k is of the order of 1 pN.nm. s (or 1 × 10^-21^kg*m*^2^*sec*^-1^). Therefore, the first term in Eq. (3) is remarkably small and can be neglected in the τ_ext_-ω relation.

The actually measured τ_ext_-ω relation can be regarded as consisting of two linear regions, and the slopes of the two linear regions are – k in the respective regions of the τ_ext_-ω relation. Then, the drag coefficient k changed roughly from 0.5 to 5 pN.nm.s when the angular speed increased [16, 19, 21]. The drag coefficient deduced from the viscous force in the mixture of water and proteins. If one takes the viscous coefficient η = 1 poise = 1 × 10^-7^ pN.s/nm^2^, then the drag coefficient [1] k_η_ = 8πηa^3^ = 0.02 pN.nm.s for a rotating sphere with a radius of 20 nm, which is a factor of ten smaller than the drag coefficient in the slow region of the τ_ext_-ω relation. In the region of high rotation speed, the drag coefficient increases by another factor of ten. The source of such a high drag coefficient might be from the bearing-like structures of L- and P-rings of BFM. The frictional interaction between the bearings and the outer membrane of the cell wall might increase the drag at a high rotational speed.

Eq. (3) can be further integrated. After integration, one gets the angle of rotation *θ_i_* of the ith impulse of the torque,

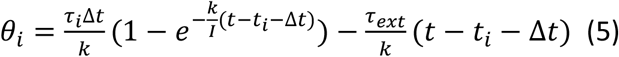

When the external torque τ_ext_ is zero, *θ_i_* exhibits a step-like behavior, which is zero before t_i_ and a constant after t_i_+Δ*t*. The exponential function in Eq. (4) is vanishingly small due to the small moment of inertia I. This is consistent with the observations that the rotation angle of BFM showed a step-like behavior when the rotation was slow. The size of the step in Eq. (4) is 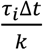 for one collision.

## Simulation experiments

I designed a simulation device to demonstrate the essence of the physical mechanism of the rotation of BFM (Fig. 2(a)). For comparison, the schematics of BFM [1, 6, 10] are shown in Fig. 2(b). In the figure, the outer membranes (Fig. 2(b)) are aligned with their counterparts (the upper thin black acrylic square plate) in the simulation device (Fig. 2(a)). A small bearing supports the rod of the rotor and functions like the L-ring of actual BFM. The inner (peptidoglycan) membrane, which accommodates the stator, is simulated with a thick square acrylic plate with a width of 10 cm. A fixed stud fixed under the lower thick plate simulates the motA of the stator of BFM, and an acrylic disk just under the stator simulates the C-ring of the rotor. The gap between the stator and the rotor is crucial; it should be a little less than the amplitude of the fluctuation of the relative positions of the stator and the rotor to ensure frequent collisions between them. The rotation of the rotor is transmitted with a rod that is free to rotate on its axis. The rod is fixed on a thin plate through a small, low-friction bearing. The asymmetry in the fluctuations of the rotor can be adjusted with the gaps (g_1_-g_4_) between the eight long-traveling nuts (Fig. 2(a)), which might correspond to the asymmetric fluctuations of the tilting angle of the L-ring of the actual BFM. For example, in Fig. 2(a), the gap gi is the smallest among the four gaps; therefore, the direction of asymmetry is from g_3_ to g_1_, which corresponds to the direction 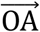 in Fig. 1. The uppermost circular black disk, which is fixed on the rod and has two white dots on it, is used to visualize the rotation. The aluminum rod with 8 mm in diameter just under the uppermost disk is designed for exerting the externally applied torque in the τ_ext_-ω relation experiments. In these experiments, three forces are exerted on the side of the aluminum rod with 120° between each of them. The torques are from three equal weights hung 30 cm from the aluminum rod to avoid the body of the electromagnetic shaker. The rotation of the rotor can be generated with a vertical electromagnetic shaker and be recorded with a digital camera for analysis.

**Fig. 2.**
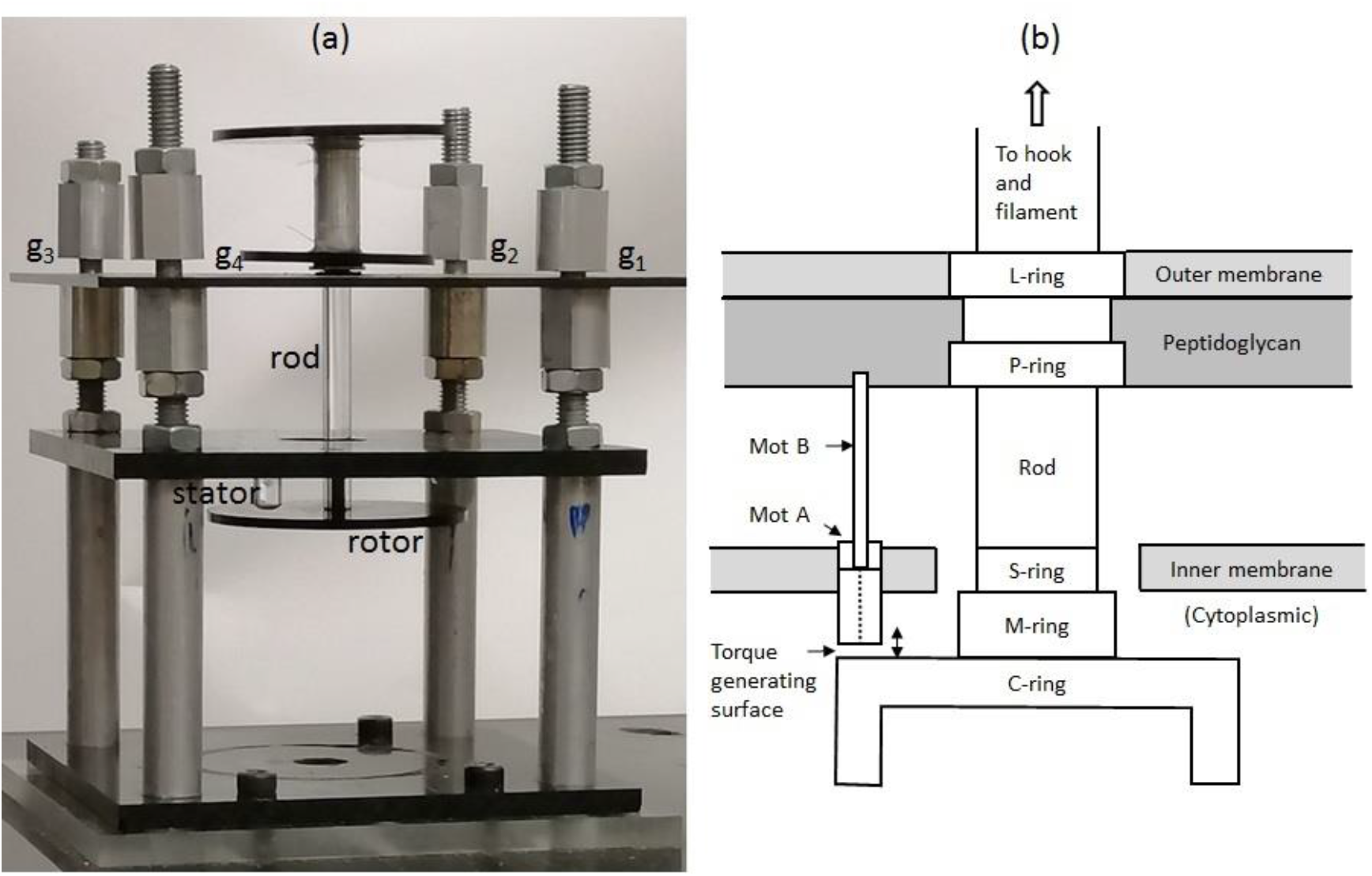
(a) Picture of the simulation device of BFM, and (b) is the schematic drawing of the structure of the actual BFM. The upper thin square plate in (a) has the function of and is aligned with the outer membrane of BFM in (b). In (a), the rod is fixed on the upper thin plate with small bearings, and g_1_, g_2_, g_3_, and g_4_ are the gaps that adjust the asymmetry of the angular fluctuations of the rod. The long screws are for adjusting the gaps. In (b), C-ring has the function of the rotor, Mot A is the part of the stator colliding with the rotor, and the surface where the stator and the rotor collide is noted as the torque-generating surface.

The net effects of the torque will be the sum of all of the generated and applied torques. The asymmetry in the fluctuations of the rotor is critical for the rotation of the rotor. If the probabilities of the occurrences of positive and negative α are equal as in the symmetric cases (for example: g_1_ = g_2_ = g_3_ = g_4_ = 3 mm), the net rotation would fluctuate around zero angle or no directional rotation. Only when steric asymmetry holds does the system have a preferred direction of rotation. A small asymmetry (α) of a fraction of a degree will be enough to generate processive directional rotation. In the proposed mechanism, the direction of rotation depends on the sign of the tilt angle; as the sign of α can be positive or negative, the rotation of BFM is bidirectional. From Eq. (1), the position of the stator and the sizes of the gaps are both essential to determine the direction of rotation. According to Eq. (1), the direction of rotation from CCW to CW (or to change the direction of tilt) can be changed in three ways by adjusting the asymmetry. The direction of rotation can be changed, in Fig. 2(a), if the position of the stator is fixed, (a) by reducing g_3_ so that g_1_ > g_3_, and g_1_ = g_2_ = g_4_, and (b) by keeping g_2_ = g_3_ = g_4_, and enlarging g_1_ so that g_1_ > g_2_, or (c) by changing the position of the stator, if the sizes of gaps are fixed. For actual BFM, the asymmetry might change its sign by rotating BFM 180° keeping the environment the same, or by switching to another active stator such that the lever arm vector (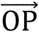 in Fig. 1) switches direction.

The images of the uppermost circular disk were recorded at 10 Hz, while the simulation device was shaken by an electromagnetic shaker at 19.7 Hz. The angles and direction of rotation of the two white spots on the uppermost circular disk were recorded.

## Results

### (a) Bidirectional rotation

It is well documented that tethered bacterial flagella change the direction of rotation abruptly and randomly [1, 2, 15]. In simulation experiments, I found that the direction of rotation could be changed easily by adjusting the conditions of the asymmetry in the fluctuations of the rotor (Eq. (1)). The time traces of the total angle that the rotor rotated was measured for various tilting angles α (Fig. 3(a)). The averaged tilting angle was changed by adjusting the size of g_1_ but the other gap sizes were maintained at 3 mm. When the simulation device was shaken vertically, the rotor rotated CCW if g_1_ was small, ceased to rotate for g_1_ = 3.5 mm, and rotated CW when g_1_ was >3.5 mm. The data indicated that the direction of rotation depended on the small asymmetry in the tilting angle, and the rotor did not rotate processively in one direction when the fluctuation in the tilting angle α was symmetric, which is consistent with the proposed mechanism. With the distance between the posts is 7 cm, the change in average tilting angle α was 0.8° for a 1-mm change in the gap size. Fig. 3 (b) shows the angular speed as a function of the g_1_ size for the time traces shown in Fig. 3(a). The angular speeds were taken as positive for CCW rotation. For g_1_ > 3 mm, the angular speeds were CW and were low compared with those of the CCW rotations. The reason might be that for the large g_1_ > 3 mm, in some occasions, the tilts of the rotor were in favor of CCW rotation, thus slowing down the CW rotation. This is what has been observed for the BFM of *E. coli* [15].

**Fig. 3.**
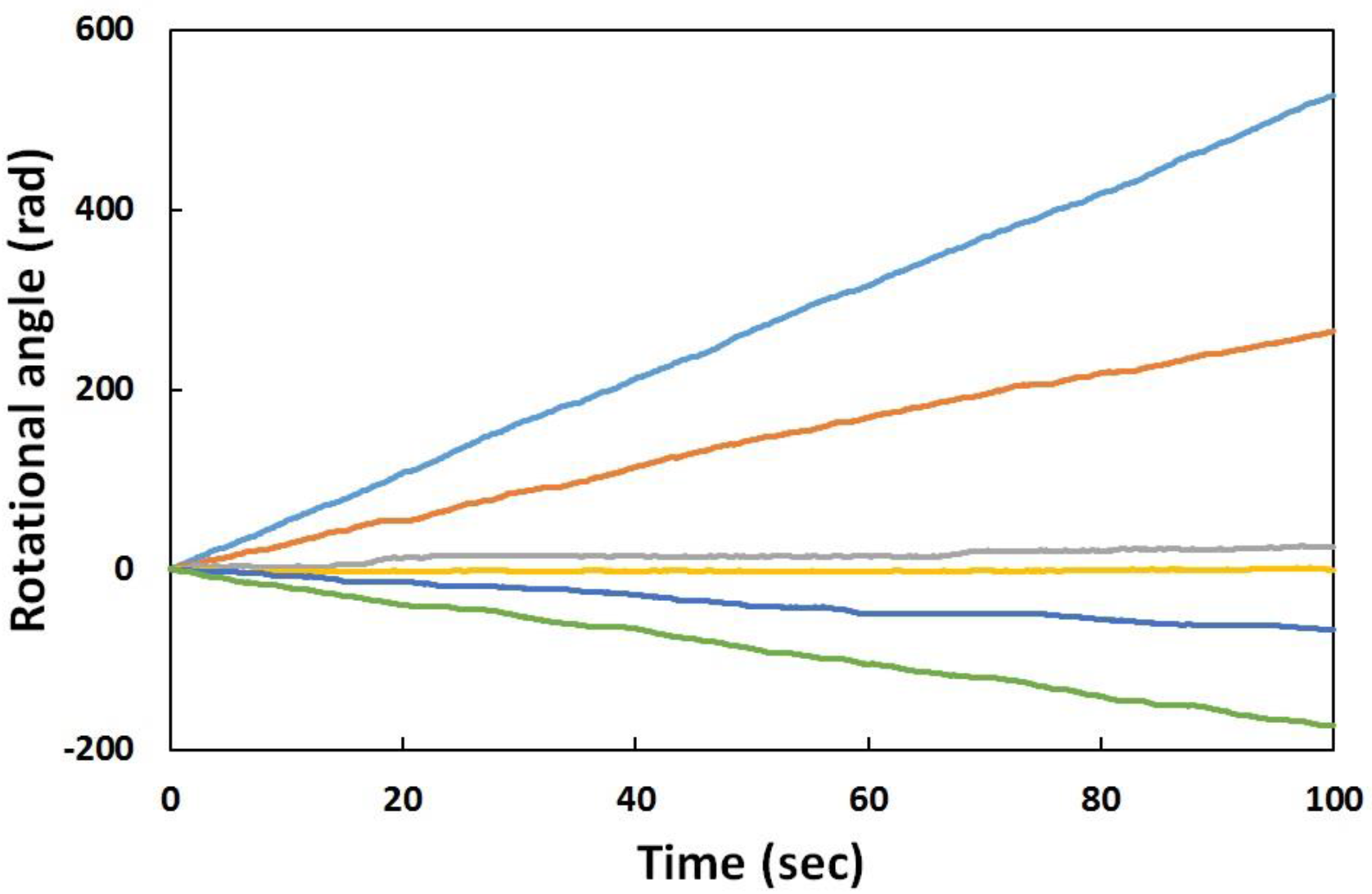
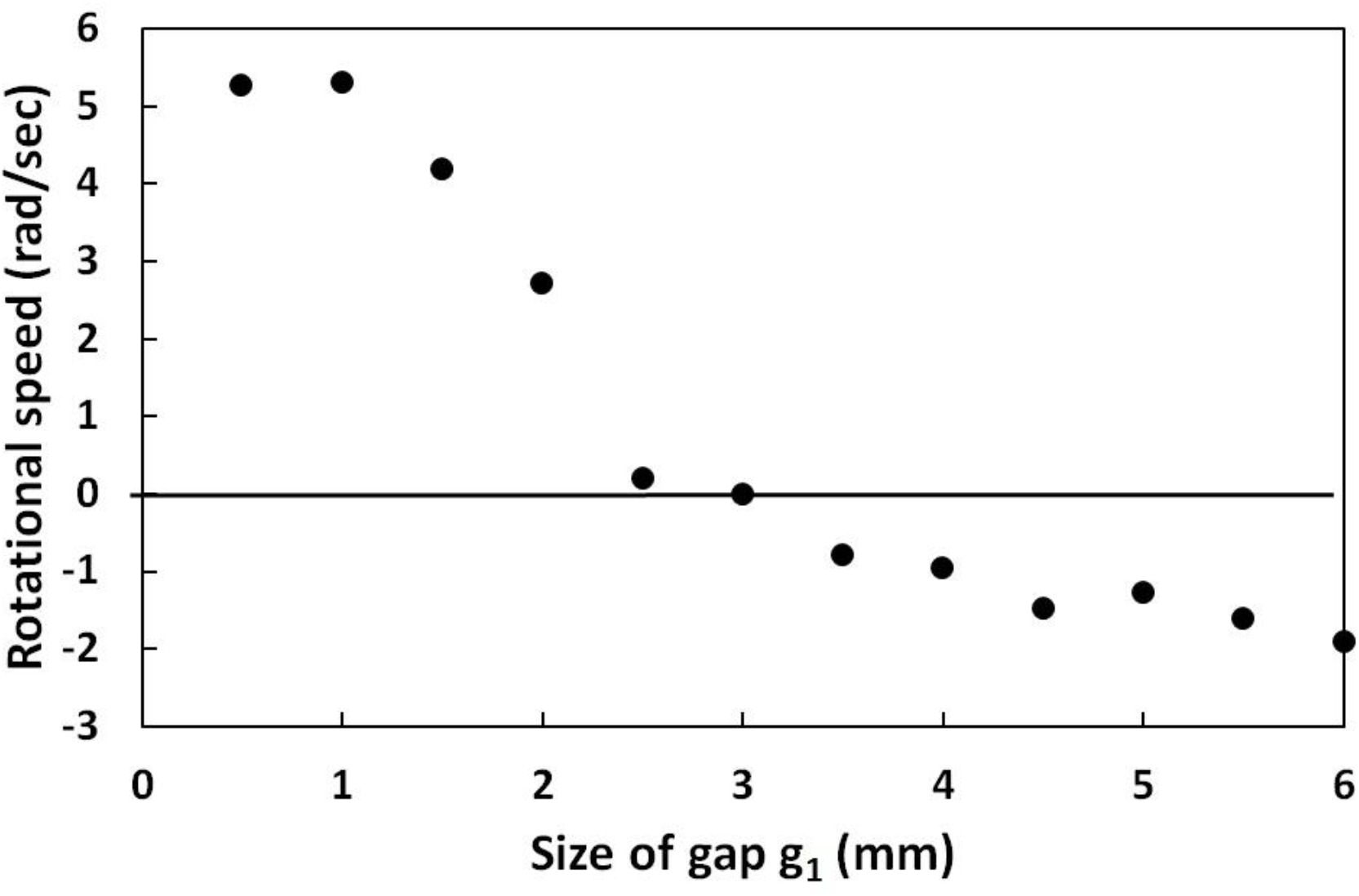
(a) Time traces of the rotational angle with different g_1_ than that in Fig. 1(a) and the other gaps are kept at 3 mm. From top down, the lines correspond to g_1_ = 1.0 mm, 2.0 mm, 2.5 mm, 3.0 mm, 3.5 mm, and 5.5 mm, respectively. (b) shows the average rotational speed as a function of the gap size g_1_. The speeds of CCW rotation are assigned as positive values. The error bars are the standard deviations calculated from five measurements. The vibrational acceleration of the shaker is set at 4 G; 1G = 9.8 m/s^2^.

### (b) τ_ext_-ω relation

The τ_ext_-ω relation is the most crucial characteristic of the rotation of BFM in model checking [1, 4, 15, 16], where the external torque was exerted on the motor in a direction against the original rotation. In high τ_ext_ region, the rotational speed is low and increases rapidly but τ_ext_ decreases. The increase in the rotational speed slows down when the torque passes a knee value. In other words, the τ_ext_-ω relation concaves downward. The asymmetry in the CCW and CW rotations shown in Fig. 3 can also be observed in the τ_ext_-ω relation in the simulation experiments [15]. As shown in Fig. 4, where the angular speeds of CCW and CW rotations are both taken as positive values, the characteristics of the τ_ext_-ω relation of the actual BFM are reproduced in the simulation experiments. Both the speed and the stall torque of the CW rotation are lower than those of the CCW rotation—consistent with the observations for the actual BFM. The simulation design preserves much of the characteristics of the actual BFM. According to the notion that the common shape of the τ_ext_-ω relation is an indication of the common mechanism for BFM [1, 5, 16], one might infer that the physical mechanism proposed in this report is one of the possible candidates of the mechanism of the rotation of BFM.

**Fig. 4.**
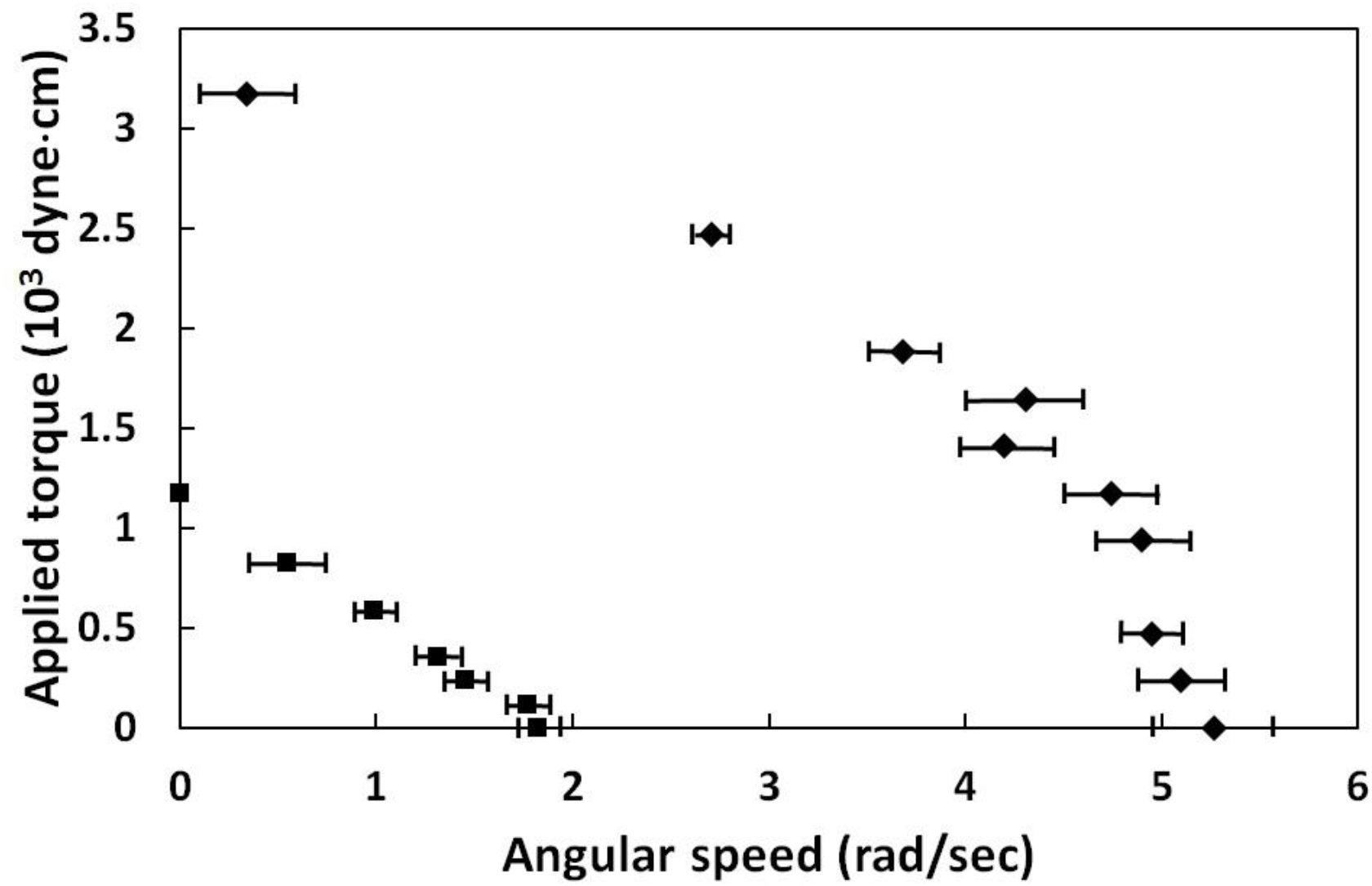
The τ_ext_-ω relations of the CCW rotation (g_1_ = 1.0 mm, diamonds) and the CW rotation (g_1_ = 5 mm, squares) for the vibrational acceleration of the shaker being set at 4 G. The error bars are the standard deviations calculated from five measurements.

### (c) The steps in rotation

In the proposed mechanism, the rotation of the rotor is generated by discrete impulsive torques. The steps in the rotation angle are genuine in the mechanical mechanism. In the simulation experiment, the discrete impulsive torques can be generated by tapping the upper plate with a finger to produce discrete collisions between the rotor and the stator. Fig. 5 shows an example of a series of steps generated by tapping. The step sizes observed here could not be compared with those observed in BFM due to different *τ_i_* and k. One characteristic of the shapes of steps in the rotation is that the steps may occur abruptly, then round off slowly to a saturation value, and wait for next impulsive torque to occur. Therefore, the edges of the steps are not extremely sharp, which can also be observed in the actual BFM experiments [17, 18]. For the actual BFM, steps were observed only when the concentration of ATP was low and the rotation was slow. Under these conditions, the occurrence of effective collisions between the stator and the rotor was rare, and the discrete steps occurred according to the mechanical mechanism.

**Fig. 5.**
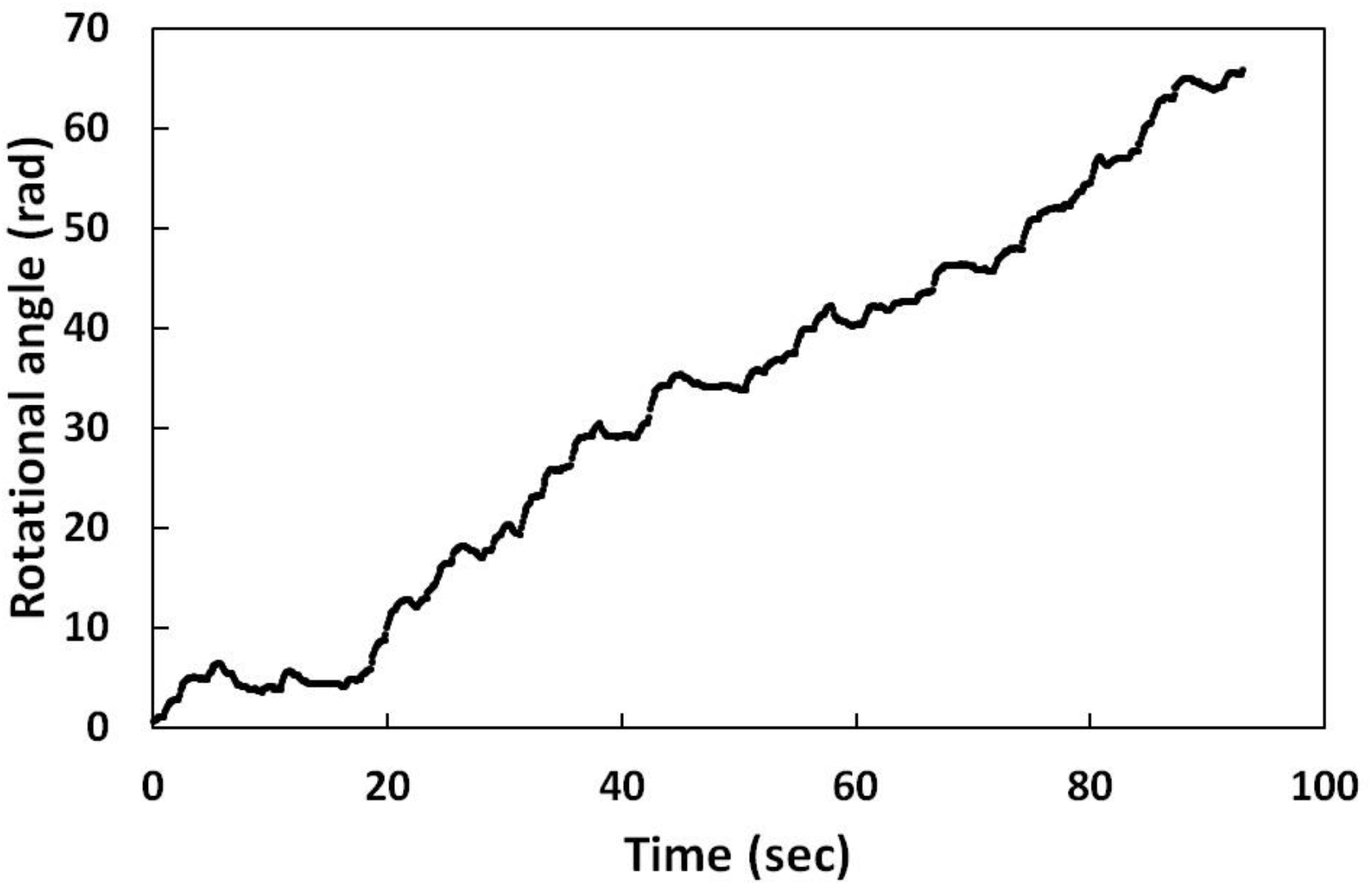
Example of time trace of the rotational steps by discrete finger tapping on the thin upper square plate of the simulation device. The edges of the steps are not as sharp as Eq. (5) predicted.

The step size of BFM can be estimated according to the result of Eq. (4), the generated torque, and the measured drag coefficient. The large drag coefficient measured from the τ_ext_-ω relation might prevent the weak collisions between the stator and rotor from producing significant rotations. The number of effective collisions might be much smaller than the total number of collisions. From Eq. (2), the average value of the generated torque τ_gen_ is 41 pN.nm per collision. However, the measured torque was between 2700 and 4600 pN.nm [1, 16], which means that N effective collisions might contribute to one step: here, N ≈ 60. Such a frequency (6×10^5^ Hz) of the effective collisions within the average correlation time of a step of 0.1 ms [6, 21] is notably low compared with the frequency of the occurrence of any collisions, which is of the order of 10^10^ Hz. According to Eq. (4), the size of the step (Δθ) is 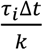 for one collision. Taking the measured drag coefficient k to be 1 pN.nm.s and the correlation time of a step Δ*t* ≈ 0.1 ms, one gets Δθ (in degrees) 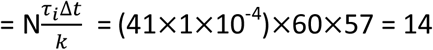, which incidentally matches the peak of the distribution of the measured step sizes [17, 18]. Steps can only be observed under unfavorable conditions for the rotation of BFM. In ordinary conditions, the rotation of BFM consists of a series of fast steps.

### (d) Dependence on the polar angle ϕ

The aforementioned results are for one stator case, and the stator situated in a direction 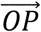, which was perpendicular to the vector 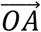. It was found that the rotational speed depended on the position of the stator P. Fig. 4 shows the rotational speed as a function of the angle ϕ, where ϕ is the angle between 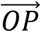 and 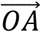 and the length of 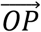 was kept the same for all of the angle ϕ. From Eq. (1), the dependence should be sinusoidal. The data shown in Fig. 6 are distorted sinusoidal dependence, probably due to the asymmetry introduced in the design of the device. The essence of the results shown in Fig. 6 is that the direction of rotation can be changed by two factors; one is the angle ϕ, which depends on the position of the stator relative to the direction of asymmetry 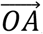, the other is the direction of asymmetry 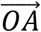, which can be adjusted by the four gaps, g_1_-g_4_. For a fixed configuration of the gaps (for example: g_1_= 1 mm, g_2_ = g_3_ = g_4_ = 3 mm) and ϕ > 180°, the direction of rotation would be CW. In this case, if one enlarges g_1_ to 5 mm, then the rotational direction would change to CCW; however, according to the same arguments as that in the first paragraph in the Results section, the rotational speed would be slower than that of the CW rotation, i.e., the speed of the CCW rotation is not necessarily larger than that of the CW rotation, as observed in Ref. 13.

**Fig. 6.**
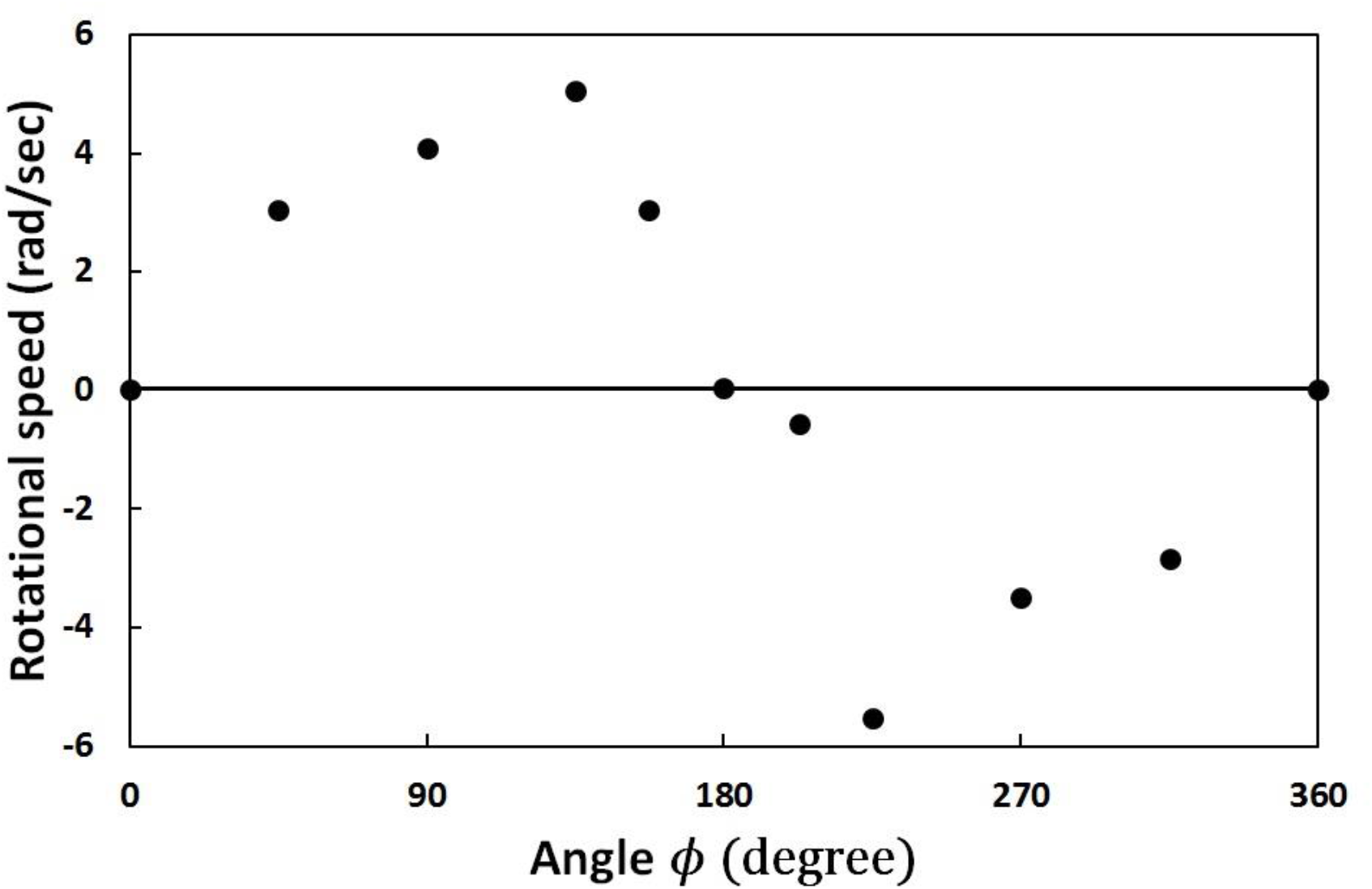
Rotational speed as a function of the polar angle *ϕ* of the position of the stator.

Another well-known observation is that some BFMs abruptly change the direction of rotation. From the results shown in Fig. 6 and the dynamics of the stators, which can be dissociated from or incorporated into the cytoplasmic membrane, I postulated that the abrupt switch of the direction of rotation might be due to the switch of the effective stators at the positions with opposite ϕs on the cytoplasmic membrane. Such a switch might occur rapidly and immediately affect the direction and magnitude of the torque generated, leading to abrupt change in the direction and the rotational speed.

### (e) Effects of multiple stators

The number M of stators involved in torque generation has been a key topic in studies on BFM [1, 19–22]. The steps observed in the resurrection experiments were regarded as an indication of the gradual increase in the number of the stators that returned to the active state. This argument was based on the assumption that the number of active stators was the only factor affecting the rotation speed of BFM. However, from Eq. (1), the torque generated, and thus the rotation speed, depends on many other factors, such as the configuration of BFM on the membrane and the positions of the stators. In the proposed mechanical mechanism, only one stator may collide with the FliG in the C-ring for any colliding (torque generation) events; in other words, only one effective stator exists at any moment. From the data shown in Fig. 6, and also from Eq. (1), the magnitude and direction of the torque generated (or the rotational speed) depend on the position of the stator. The contributions at uncorrelated moments, which might be CW or CCW, might add up without coordination. Therefore, the rotation speed might not strongly depend on the number of the stators in the cytoplasmic membrane [4, 6].

For multiple stators, the dependence of the rotation speed (or torque) on the number M of stators may be different for different distributions of the stators in the cytoplasmic membrane in the proposed mechanical mechanism. Fig. 7(a) shows the dependence of the rotation speed on the number of stators for two different stator arrangements. The full dots are for the case that all of the stators are arranged in the same half of a circle (relative to the direction of asymmetry 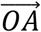) in the plate simulating the cytoplasmic membrane with ϕ < 180° and CCW rotation; CW makes no contribution in this arrangement. Due to the limited space, only seven stators are put on the plate. The rotation speed does not increase linearly with M and actually decreases a little with a large M. This is probably because, occasionally, the rotor collides with the stators situated at the place with ϕ deviating largely from 90°, which, as per Eq. (1), contribute less to rotation speed. The full triangles in Fig. 7(a) are for the case that the stators are distributed evenly (one for every 45°) in a circle centered at O in the plate simulating the cytoplasmic membrane. The rotation speed (CCW) decreases rapidly for large M. The reason for the decrease might be that stators with ϕ ≥ 180°, as per Eq. (1), contribute negative (CW) torque to the rotation; thus, the collisions of the rotor with such stators reduce the rotation speed when M and ϕ become large. In the case of seven evenly distributed stators, the asymmetry is lost and any angle ϕ is equally probable to occur in the torque-generating collisions. Therefore, the generated torques cancel each other, leading to the rotor fluctuating around the initial angle with no net rotation.

**Fig. 7.**
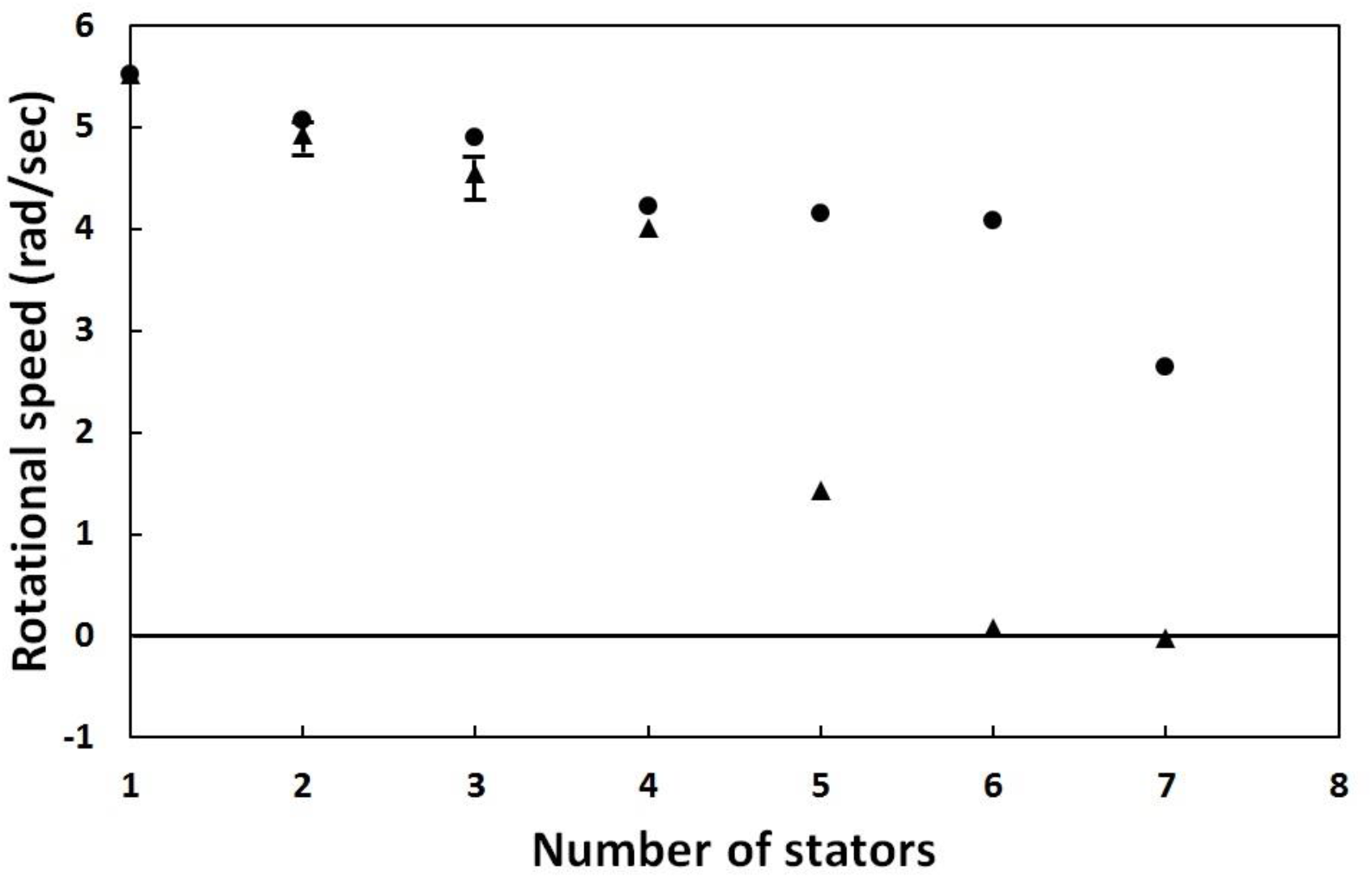
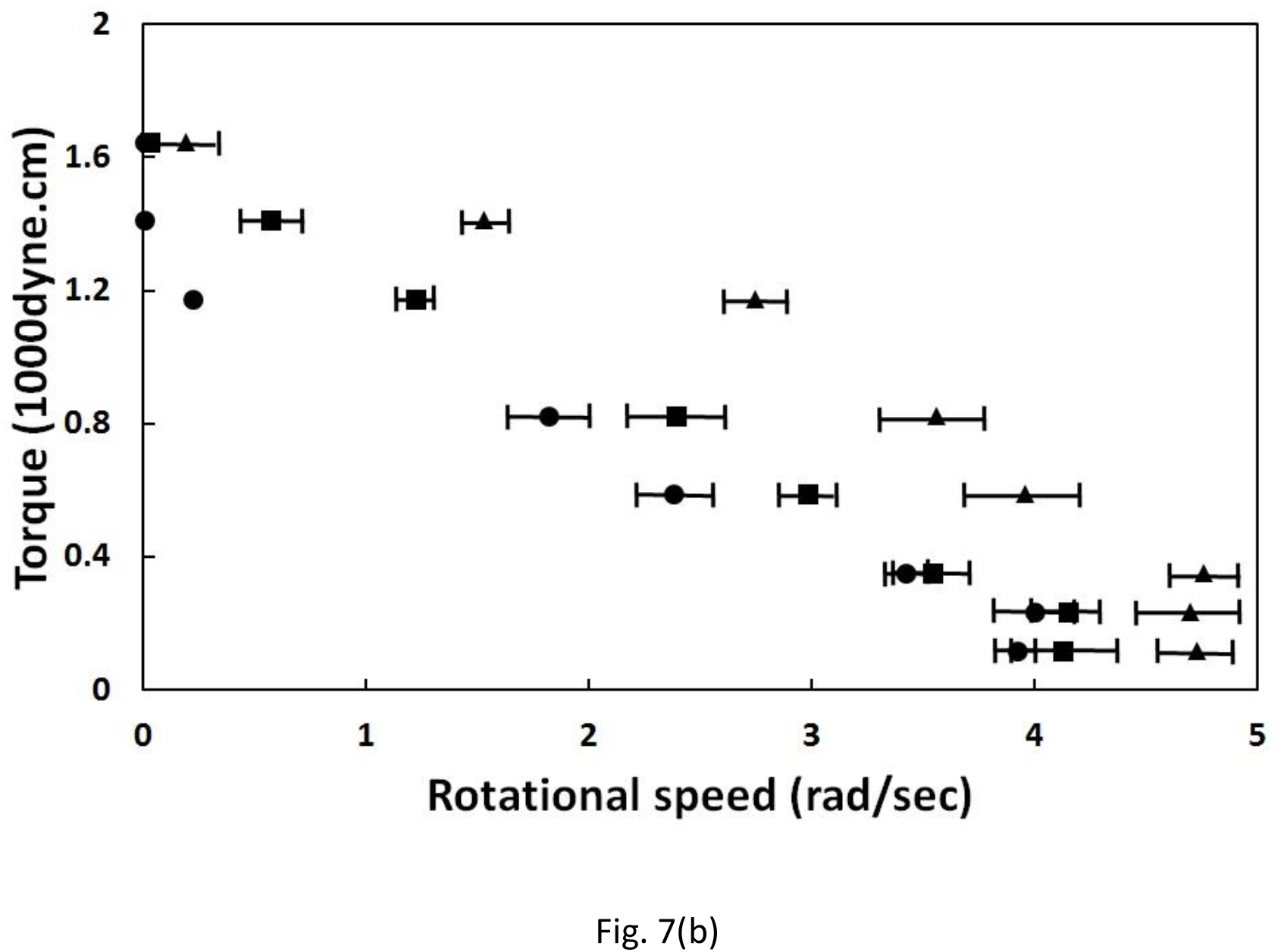
(a) Rotational speeds as functions of the numbers of stators for two different stator arrangements. The full circles are for the stators arranged in the same half-circle in reference to the line of asymmetry 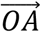. The full triangles are for the stators set every 45° apart where the angles are measured in reference to 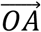. (b) The τ_ext_-ω relations of one (full triangles), three (full squares), and five (full circles) stators arranged in the same half-circle in reference to the line of asymmetry 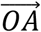.

An increase in the number M of the stators did not increase in the power output in the simulation experiments either. Fig. 7(b) shows the τ_ext_-ω relations with one, three, and five stators deposited in the same half-circle relative to the direction of asymmetry 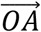 in the plate simulating the cytoplasmic membrane. When adding stators to the plate, the ϕs of the individual stators are different and are kept <180°, so that the torques generated with the stators are positive and contribute to CCW rotation. As shown in Fig. 7(b), the increase in the number of stators decreases not only the stall torque but also the maximum rotational speed.

## Discussion

The flagellar rotation is under continuous viscous drag and in the overdamping regime. A reliable and strong source of torque is required for the processive rotation of the flagella. Finding the origin of the generation of torque is a purely mechanical problem, and a clear source of the force and the lever arm of the force to generate the torque must be specified. In the mechano-electrochemical mechanism [5,6] of the rotation, the source of the required force was proposed to be from the conformation changes of the stators and the steric interaction potential between the stators and the rotor. However, such an interaction potential may not be the dominant source of the force of interaction between the stator and the rotor. The impulsive force due to the physical collisions between the stator and the rotor may be much stronger than the force from the continuous interaction potential. Therefore, I propose a purely mechanical explanation for BFM, namely that the torque generated from the impulsive force causes the flagellar rotation. The collisions between the stator and the rotor originate from the thermal motions of the two parties, which produce the impulsive force. The asymmetric fluctuations of the azimuthal angle of the rotor and the off-center point of collision generate the averaged directional torque (Eq. (1)).

The rotational speed depends linearly on the ambient temperature, and the observed slope of the linear dependence is 5.4 Hz/°C [26]. From Eq. (3) for steady rotation, 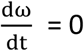 and *τ_ext_* = 0, *kω* = *τ_gen_*. From Eq. (2) and considering the number of correlated collisions, ω = N(R/kd)k_B_Tsinϕsinα, where N ≈ 60, R = 20 nm, d ≈ 0.2 nm, the drag coefficient k ≈ 1 pN.Nm.s, the Boltzmann constant k_B_ = 1.38×10^-23^ J/°k = 1.38×10^-2^ pN.nm/°k, and sinϕsinα ≈ 0.1, then ω ≈ 8.3T. The measured frequency f = ω/2π = 1.3T. The slope of the linear dependence of f on temperature T is estimated to be 1.3 Hz/°C, which is of the same order of magnitude of the measured slope. Considering that the estimation is only a rough one, the result supports the proposed physical mechanism of the torque generation of BFM.

The applicability of a proposed mechanism can be proven by many methods—for example, model calculations, computer simulations, or simulation with a real device. If the methods are to be eligible to prove a mechanism, the individual proving method must contain the essential characteristics for both the structure and the physical principles involved and can be used to obtain the observed results of the actual system. In this study, I designed a device that has the key structures of BFM as described in the section of the design of the simulation device (Fig. 2), and the only interactions between the stators and the rotor are physical collisions. The simulation experiments reproduced most of the observations of BFM, such as bidirectional rotation, τ_ext_-ω relation of CCW and CW rotations, and steps in the time trace of the angle of rotation. Therefore, the contribution from the mechanical mechanism cannot be overlooked. In the mechanism, the torque might be generated from the random collisions of the stator with the rotor. Such a generation of rotation from random motion does not violate the second law of thermodynamics. Because BFM is not an isolated energy system, the static electric energy released due to positive ion flow through the potential difference across the bacterial membrane to BFM enhances the random motion of the rotor and partially powers the rotation.

## Summary

A physical mechanism that the torque requires for the rotation of the rotor of BFM is generated from the impulsive force due to random collisions between the stator and the rotor is proposed for the first time. The CCW or CW direction of rotation is determined by the asymmetry in the fluctuations of the tilt angle of the rod. The drag coefficient derived from the τ_ext_-ω relation is too large to be accounted for by the viscous force from the fluid environment. Instead, drag from the interaction between the L-ring and the outer membrane might be the origin of the observed drag coefficient. The expressions of the torque and the step size can be derived by directly solving a Langevin equation of motion (Eq. (3)).

The order-of-magnitude estimations of the torque, the step size, and the temperature dependence of the rotational speed are consistent with previous experimental observations. The estimations are based on the experimental data of BFM and have only two adjustable parameters—the number N of the effective collisions per step and the average tilt angle α—and the choices of the two parameters are closely related. The abrupt change in the direction of rotation might be attributed to the abrupt switch of the stators on the opposite side of the direction of asymmetry (opposite in ϕ).

The energy released by the ions flowing down the electric potential difference across the flagellar membrane is most probably dissipated to the bulk of BFM along the path of flow. The released energy might power the rotation of BFM indirectly by enhancing the random motions of the stator and the rotor. The applicability of this proposed mechanical mechanism was demonstrated using a simulation device processing the structural characteristics of BFM. Numerous observations for the actual BFM, such as bidirectional rotation and torque-speed relations of the CW and CCW rotations, were reproduced in simulation experiments.

## Acknowledgment

The work is supported by the physics department of the National Tsing-Hua University.

